# An uncertainty-based interpretable deep learning framework for breast cancer outcomes prediction

**DOI:** 10.1101/2022.08.25.505346

**Authors:** Hua Chai, Siyin Lin, Minfan He, Yuedong Yang, Yongzhong OuYang, Huiying Zhao

## Abstract

Accurate prediction of breast cancer outcomes is important for selecting appropriate treatment, which can prolong the survival period of the patients and improve the life quality. Recently, different deep learning-based methods are carefully designed for cancer outcomes prediction. However, the applications of these methods are still challenging due to the model interpretability. In this study, we proposed a novel multi-task deep neural network UISNet to interpret the feature importance of the prediction model by an uncertainty-based integrated gradients algorithm. Additionally, UISNet improves the prediction accuracy by introducing the prior biological pathway knowledge and utilizing the patients’ heterogeneity information. By applications to seven breast cancer public datasets, the method was shown to outperform state-of-the-art methods by achieving a 5.79% higher C-index value on average. For the identified genes based on the interpretable model, 11 out of the top 20 genes have been proved to be associated with breast cancer by literature review. The comprehensive tests indicated that our proposed method is accurate and robust to predict breast cancer outcomes, and is an effective way to identify the prognosis-related genes. The method codes are available at: https://github.com/chh171/UISNet.

## 1. Introduction

According to the report on new cases of 36 common cancers in 2020, breast cancer is the most common malignant tumor that accounted for 2,261,419 (11.7%) patients over the world^[1]^. The breast cancer patients’ outcomes were observed significantly different with the same treatment, which contributes the most to hindering the effectiveness of the therapies. Accurate prediction of the breast cancer outcomes is important for patients’ follow-up treatment, which can prolong the patients’ survival period and improve their quality of life^[2]^. With the development of molecular sequencing techniques, more and more high-dimensional genomic data were used to evaluate the cancer outcomes, for supporting clinical decision making^[3]^.

The most widely used method used to evaluate patients’ risks is the Cox proportional hazard model^[4]^. The method analyzes the influence of different factors on cancers by calculating the survival ratio of patients without knowing the survival distribution of the patients. Besides, Wang *et al*. designed the random survival forests (RSF) for predicting cancer outcomes by utilizing the bootstrapping strategy^[5]^. However, these methods didn’t perform well with the high-dimensional gene expression data^[6]^. To solve this problem, different dimension reduction technologies were added in the algorithm. Lin used features extracted by principal component analysis (PCA) in the Cox method to predict disease prognosis^[7]^. Considering PCA performed poorly in the high-dimensional nonlinear space, Cai selected kernel PCA instead of PCA to generate the compressed features with different kernel functions, and put them into the Cox framework for patient’s risk prediction^[8]^. Another way to solve the computational challenge caused by high-dimensional features is to add the regularization part to the Cox model. Boulesteix combined the adaptive Lasso regularization and the Cox regression method (IPW-lasso) to estimate the cancer prognosis^[9]^. The method minimized the likelihood function by an L1-norm regularization constant, to shrink the coefficients of the features. In addition, the survival support vector machines (SSVM) proposed by Evers improved the performance of the cancer outcomes prediction by using a sparse kernel function. However, it is a complex process for choosing the suitable kernel function and hyper-parameters. Recently, Liu *et al*. designed an integrated learning method EXSA based on the XGBoost framework, to evaluate the cancer prognosis. The results show it outperformed other traditional machine learning methods^[10]^.

In recent years, Deep learning (DL) technologies show advantages in dealing with high-dimensional nonlinear features. Deep_surv proposed by Katzman was designed to estimate cancer outcomes by combining the deep neural network (DNN) and the proportional hazard loss function^[11]^. Chaudhary used the Autoencoder to reconstruct the high-dimensional expression features, and the generated features were used in Cox proportional hazards model for liver cancer survival analysis^[12]^. Based on this method, Yang *et al*. proposed DCAP by using the denoising Autoencoder to improve the ability of the DNN against the data noise^[13]^. Nonetheless, the separation of feature extraction and risk prediction hindered the convenience of the method. To solve this problem, an end-to-end DNN framework was proposed by combing the loss of risk prediction and data recovery loss^[14]^. On the basis of these studies, for speeding up the convergence of deep neural networks and reducing the overfitting in model training, Qiu designed a meta-learning-based network for cancer outcomes prediction^[15]^.

Though the DL-based methods achieved better results in cancer outcomes prediction, the application of these methods is still limited by the lack of the model interpretability. Based on the interpretability of the prediction model, identifying the cancer-related biomarkers has important implications for auxiliary medical decision making and target-drug development. The widely used solution to solve this problem in DL-based methods is differential expression analysis (DEA). Nevertheless, when the average expression of the features is low, the log-fold change values computed in DEA are disproportionately affected by the noise^[16]^. Hence, Hao proposed an interpretable DNN framework for cancer survival analysis, by calculating the gradients in the neural network^[17]^. Whereas, this approach may lead to gradient saturation, making the neural network difficult to identify some important features^[18]^. In another way, Zhao *et al*. proposed DeepOmix for cancer prognosis prediction^[19]^. According to the predicted risks, DeepOmix performed the Kolmogorov-Smirnov test to identify the prognosis-related genes. However, the features selected by DeepOmix may not necessarily be related to the DNN output. The prediction model may still predict the cancer outcomes accurately after removing these genes, whereas only using these features did not always get satisfactory prediction results.

To address these problems, we propose an uncertainty-based interpretable deep semi-supervised network (UISNet) for breast cancer outcomes prediction. The main contributions of UISNet are given as follows:

1. An uncertainty-based algorithm that combines the Monte Carlo dropout and the integrated gradients is designed to improve the reliability of the interpretable results.
2. By introducing the prior biological pathway information as a sparse layer, UISNet interprets the feature importance of the prediction model while efficiently processing high-dimensional gene expression data.
3. UISNet gets more useful information for cancer outcomes prediction by integrating the patients’ heterogeneity clustering, cancer outcomes evaluation, and dimension reduction tasks into a unified framework. These tasks are simultaneously optimized by the shared representations in the neural network.
4. We made a comprehensive analysis by collecting seven breast cancer datasets from TCAG and GEO databases. The results proved that UISNet is accurate and robust to predict breast cancer outcomes, and is an effective way to identify the prognosis-related genes.

The remainder of this article is organized as follows. The details about UISNet are introduced in Section 2. Section 3 shows the experiment result and the biological analysis. At last, we give the conclusion and discussion in Section 4.

## 2. Materials and Methods

### 2.1 Datasets

Seven breast cancer datasets from GEO (https://www.ncbi.nlm.nih.gov) and TCGA (https://www.cancer.gov/about-nci/organization/ccg/research/structural-genomics/tcga) were used for method evaluation in this study. The common 4767 gene features in these datasets were normalized by log transformation, and the batch effect was removed by using the “*limma*” package^[20]^. The number of uncensored samples (Uncen), total samples, and the censoring rate (CR) of the used datasets are given in Table 1.

**Table 1.** The statistical information of used cancer data in TCGA

| Dataset | Uncen | Total | CR (%) |
| --- | --- | --- | --- |
| BRCA | 79 | 609 | 87.03 |
| GSE2990<sup>[21]</sup> | 48 | 125 | 61.60 |
| GSE9195<sup>[22]</sup> | 12 | 77 | 84.42 |
| GSE11121<sup>[23]</sup> | 46 | 200 | 77.00 |
| GSE17705<sup>[24]</sup> | 71 | 298 | 76.17 |
| GSE19615<sup>[25]</sup> | 101 | 115 | 12.17 |
| GSE25066<sup>[26]</sup> | 111 | 508 | 78.15 |
| Total | 468 | 1932 | 75.78 |

### 2.2 The architecture of the UISNet

As shown in Fig 1, the high-dimensional gene expression data are given in the input layer, and the prior biological pathway information is introduced in the sparse layer. UISNet can learn meaningful information by incorporating the prior biological knowledge in the sparse layer (e.g., KEGG and Reactome gene connection pathways). The knowledge of the learned relationships between genes and function pathways is used to form sparse connections between the input layer and the sparse layer instead of the fully-connections.

**Fig1.**
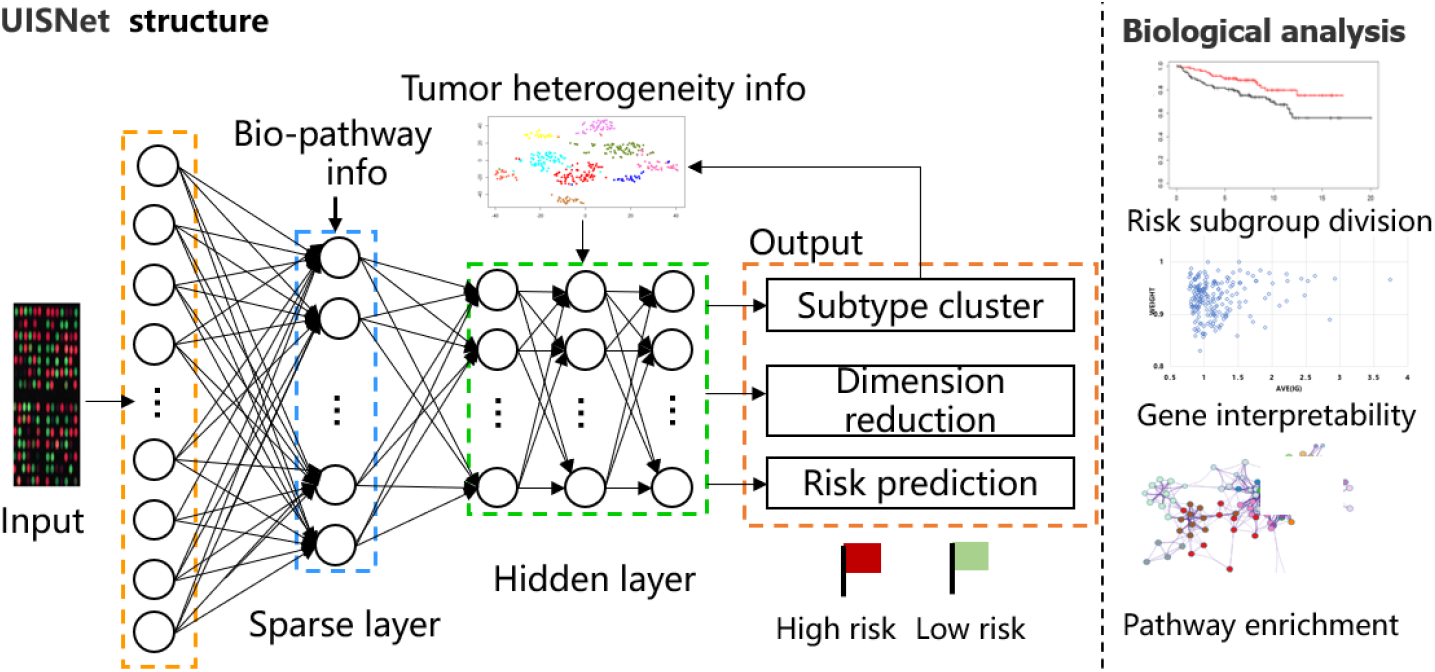
The architecture of UISNet for breast cancer outcomes evaluation. The prior biological pathway information is introduced in the sparse layer, and UISNet predicts breast cancer outcomes by integrating the patients’ heterogeneity clustering, dimension reduction, and cancer outcomes prediction tasks into a unified framework.

The binary bi-adjacency matrix is constructed to represent the sparse connections between the genes and functional pathways as in [17]. Supposing *p* gene features and the prior information including *q* pathways from KEGG and Reactome databases are given in the neural network. The binary bi-adjacency matrix can be expressed as: *Aϵ*𝔹^*q*×*p*^, the element *a*_*ij*_ in *A* is given as:

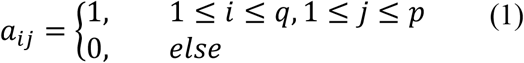

The node values *h* in the UISNet framework are computer as:

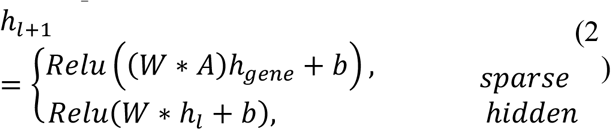

where *Relu* is the nonlinear activation function, *h*_*gene*_ represents the gene expression values, *h*_*l*_ is the output in layer *l, W* is the weight matrix, and *b* is the bias, respectively.

Supposing *X* = (*x*_1_, *x*_2_, … *x*_*p*_) represents the gene expression of the breast cancer patients, *Z* is the low-dimensional representation of *X* in the last hidden layer. The dimension reduction task is used to get high-quality compressed features in the hidden layer. Similar as the encoder-decoder structure, supposing *E* is the encoder function and *D* is the decoder function, the compressed *Z* is written as: *Z=E(X)*, and the recovered *X’* can be expressed: *X’=D(Z)*. The loss of the dimension reduction task is written as:

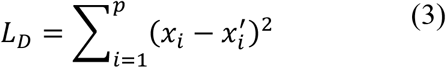

The subtype cluster task is designed to extract the information of the breast cancer patients’ heterogeneity. In the last hidden layer, the feature matrix is formed by integrating the produced *Z* and the cluster labels *L*. The cluster task loss in UISNet are defined as KL divergence between the two distributions *S* and *T* ^[27]^:

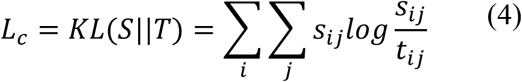

where *t*_*ij*_ describes the similarity between the cluster center *μ*_*j*_ and embedded point *z*_*j*_ by Student’s t-distribution:

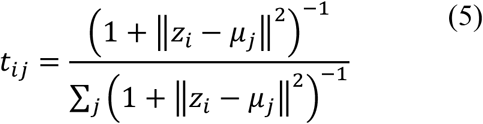

*s*_*ij*_ is the target distribution:

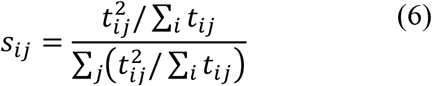

The initial cluster labels *L* of the patients are calculated by k-means. The number of the cluster *k* is chosen in [2, 3, 4] with the largest silhouette score. For ensuring the accuracy of the cluster task in the UISNet framework, the computed labels are updated in each epoch with the program running.

The risk prediction task is used to evaluate breast cancer prognosis. Supposing *S*(*t*) = *Pr* (*T* > *t*) is the survival probability that the patient will survive before time *t*. The time interval *T* is the time elapsed between data collection and the patient’s last contact (end of the experiment/ patient death). The risk function at *t* can be given:

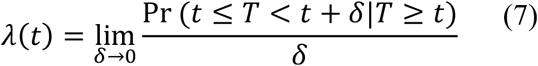

The loss function of the prognosis evaluation task can be expressed as[11]:

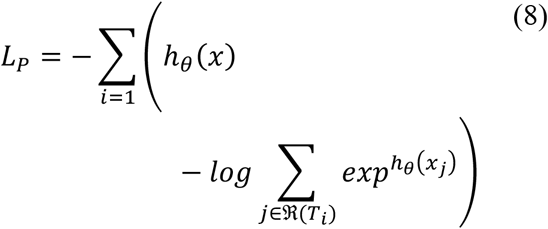

where UISNet updates *h*(*x*) by the weight *θ*, and ℜ(*T*_*i*_) represents the risk set of breast cancer patients still alive at the time point *T*_*i*_.

By integrating the dimension reduction, patients’ heterogeneity clustering, and cancer prognosis evaluation tasks into a unified framework, the total loss of UISNet can be given:

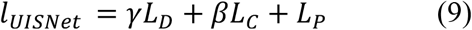

The *γ* and *β* can balance the importance of these tasks, which can be seen as the hyper-parameters chosen by the cross-validation. In this study, the value of *γ* was set 1, and *β* was set 10.

### 2.3 The uncertainty-based integrated gradients algorithm

In [17], the gradients of the output *y* with respect to the input *x* (*W* = ∂*y*/∂*x*) are used to quantify the importance of each gene to the cancer prognosis. However, computing the gradients in the deep neural network has been shown to result in gradient saturation. To interpret the feature importance of the prediction model, UISNet uses the Gauss-Legendre quadrature to approximate the integral of gradients, after calculating the gradients of the input *x* across different scales against the baseline *x*_*i*_ (zero-scaled):

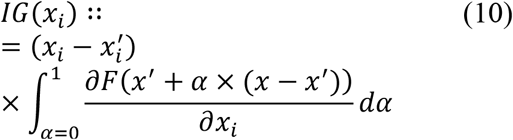

Nevertheless, the evaluation results given by the IG algorithm are not always reliable. Calculating the uncertainty of the predictions can judge the reliability of the results. Bayesian neural networks have been designed to quantify the uncertainty of results, but due to the large amount of computation, Monte Carlo dropout and Gaussian distribution models are often used as approximate solutions for Bayesian neural networks. Compared to Gaussian distribution models, Monte Carlo dropout can better approximate the approximation to Bayesian neural networks, by using the dropout as one regularization term for calculating the uncertainty of the results^[28]^. The objective function using L2 regularization in Monte Carlo dropout can be expressed as:

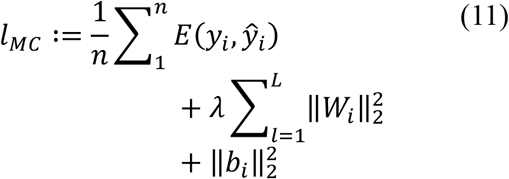

where *L* is the number of layers in the deep neural network, *y*_*i*_ and 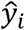 are the target and the output of the network, respectively. Following^[29]^, we trained UISNet with different dropout settings at *T* inference times as the Monte Carlo dropout strategy. Supposing the *lgx*_*i*_ represents the IG importance of the *i*th node, the uncertainty of the *i*th features is designed as:

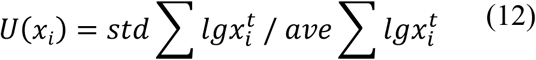

where *std*(*) and *ave*(*) are the standard deviation and average values of the *lgx*_*i*_ a t *T* inference times. The importance weight of the gene feature in the network is expressed as:

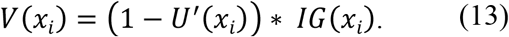

where *U’* (*x*_*i*_) is the adjusted uncertainty value of *U*(*x*_*i*_) after log transformation and min-max normalization. The larger the *V*(*x*_*i*_) value of the feature, the more important the feature is. In summary, the algorithm of UISNet is given in following:

#### Algorithm of *UISNet*

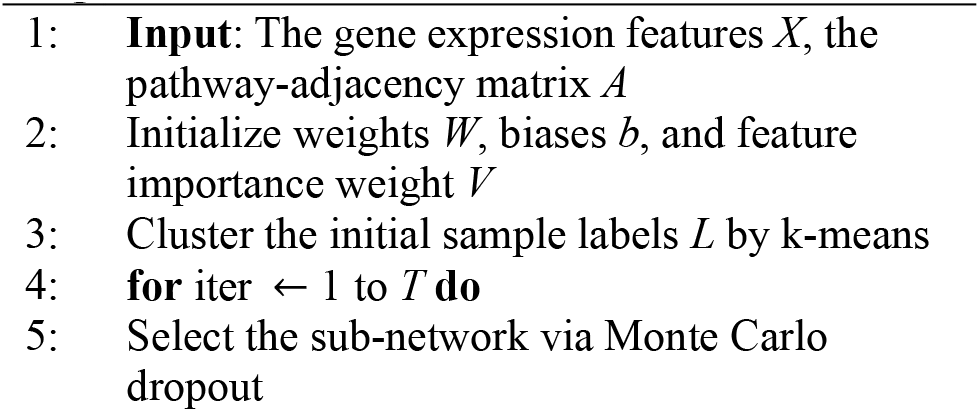
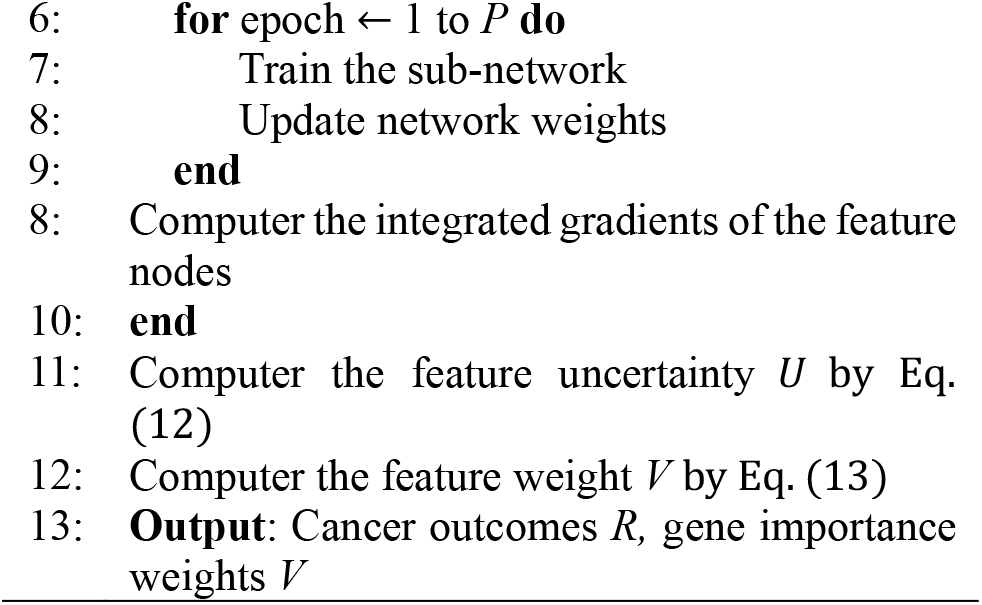

### 2.4 Performance evaluations and parameters selection

In this study, the cancer outcome prediction performances are compared through the C-index and |log10(p)| values. The C-index (CI) value is the fraction of all pairs of patients whose predicted outcomes are correctly ordered based on Harrell’s C statistics^[30]^. The higher |log10(P)| value indicates more significant differences in survival between the patient subgroups divided based on the predicted risks.

The parameter list in this study was given in below: The number of nodes in hidden layer 1 was set 1000, and the number of nodes in hidden layer 2 was set 500. The number of nodes *Z* in hidden layer 3 was selected in [50, 20, 10]. The learning rate (LR) was set to [1e-6,1e-7,1e-8], and the max epoch in the neural network was set to 2000. The optimal parameters were selected by the 5-fold CV.

## 3. Result

## Funding

This work was funded by the Key Special Project of the Department of Education of Guangdong Province (2021ZDZX2060), the National Science Foundation of China (No. 82002587), and Jihua laboratory scienctific project (X210101UZ210).

## Notes

### Competing Interest Statement

The authors have declared no competing interest.

